# Small change, big difference: A novel praziquantel derivative P96 with multistage antischistosomal activity for chemotherapy of schistosomiasis japonica

**DOI:** 10.1101/2023.03.06.531249

**Authors:** Jing Xu, Lan-Lan Dong, Huan Sun, Ping Huang, Run-Ze Zhang, Xin-Yi Wang, De-Qun Sun, Chao-Ming Xia

## Abstract

**Background:** Praziquantel (PZQ) has been the first line antischistosomal drug for all species of *Schistosoma*, and the only available drug for schistosomiasis japonica, without any alternative drugs since the 1980s. However, PZQ cannot prevent reinfection, and cannot cure schistosomiasis thoroughly because of its poor activity against juvenile schistosomes. In addition, reliance on a single drug is extremely dangerous, the development and spread of resistance to PZQ is becoming a great concern. Therefore, development of novel drug candidates for treatment and control of schistosomiasis is urgently needed.

**Methodologys/Principal findings:** A novel PZQ derivative P96 was synthesized with the substitution of cyclohexyl by cyclopentyl. We investigated the *in vitro* and *in vivo* activities of our drug candidate P96 against different developmental stages of *S. japonicum*. Parasitological studies and scanning electron microscopy were used to study the primary action characteristics of P96 *in vitro*. Both mouse and rabbit models were employed to evaluate schistosomicidal efficacy of P96 *in vivo*. Besides calculation of worm reduction rate and egg reduction rate, quantitative real-time PCR was used to evaluate the *in vivo* antischistosomal activity of P96 at molecular level. *In vitro*, after 24h exposure, P96 demonstrated the highest activities against both juvenile and adult worm of *S. japonicum* in comparison to PZQ. The antischistosomal efficacy was concentration-dependent, with P96 at 50μM demonstrating the most evident schistosomicidal effect. Scanning electron microscopy demonstrated that P96 caused more severe damages to schistosomula and adult worm tegument compared to PZQ. *In vivo*, our results showed that P96 was effective against *S. japonicum* at all developmental stages. Notably, its efficacy against young stage worms was significantly improved compared to PZQ. Moreover, P96 retained the high activity comparable to PZQ against the adult worm of *S. japonicum*.

**Conclusions:** P96 is a promising drug candidate for chemotherapy of schistosomiasis japonica, which has broad spectrum of action against various developmental stage, potentially addressing the deficiency of PZQ. It might be promoted as a drug candidate for use either alone or in combination with PZQ for the treatment of schistosomiasis.

**Author Summary:** Schistosomiasis is one of the neglected tropical diseases caused by infection of *Schistosoma spp*. Currently, in the absence of effective vaccines for schistosomiasis, PZQ is the first line drug chosen for the treatment and control of schistosomiasis in most developing countries. However, after long-term and large-scale administration of PZQ, drug-resistance has been a great concern. Therefore, there is a need for new therapies. In this study, with the aim of preventing the formation of less effective metabolite 4-*trans*-cyclohexanol, a novel PZQ derivative, P96, is synthesized with the cyclohexyl group substituted by cyclopentyl group. It is this small modification that gives us a big surprise. *In vitro*, all the biological assessments, including viability reduction rate and morphological properties by scanning electron microscopy, demonstrate that P96 has superior anti-schistosomula activity compared to PZQ, and retains similar or even higher anti-adult *S. japonicum* activity to PZQ. The antischistosomal effect of P96 is dose-dependent. *In vivo*, P96 displays high efficacy against all developmental stages of *S. japonicum*, with significantly improved efficacy against young stage worms compared to PZQ. Furthermore, the quantitative detection results of specific circulatory SjR2 DNA prove that P96 has similar activity to PZQ against adult schistosome at molecular level in rabbit sera with infection of schistosomiasis. In conclusion, the novel PZQ derivative, P96 is a promising drug candidate for chemotherapy of schistosomiasis, potentially addressing the deficiency of PZQ, and might be promoted for use either alone or in combination with PZQ for treatment and control of schistosomiasis.

## Introduction

Schistosomiasis is a relatively neglected tropical disease caused by blood flukes of the genus *Schistosoma* which afflicts more than 250 million people worldwide [1]. Globally, schistosomiasis is endemic in 78 countries, and nearly 800 million people are at risk of being infected [1–3]. Six geographically distinct species of *Schistosoma*, including *S. mansoni, S. haematobium, S. japonicum, S. intercalatum, S. mekongi, S. guineensis*, are responsible for infections in humans, resulting in significant morbidity and attributing to over 200,000 deaths per year [2–4]. The disability-adjusted life years (DALYs) caused by schistosomiasis ranges from 1.9 million to 70million according to different estimates [2–6].

To date, no efficacious schistosomal vaccine for human is available, and praziquantel (PZQ) remains the solely available drug for the treatment and control of schistosomiasis [6]. Despite its efficacy against adult worms of all schistosome species infecting humans, PZQ does not kill developing schistosomes, and cannot prevent reinfection, which is clearly the exclusive reason for the persistence of schistosomiasis [7–8]. In addition, after long-term and large-scale of mass drug administration campaigns, PZQ-resistance has been a constant concern [2–8]. In fact, PZQ-resistant isolates of *S. mansoni* have been firstly demonstrated by Fallon and Doenhoff in 1994 [9]. In 1995, the first case of acquired resistance to PZQ was recorded in Senegal [10]. Although the resistance of *S. japonicum* to PZQ has not been reported, the therapeutic dose in mainland China has increased from one 40 mg/kg dose to its current level of two 60 mg/kg doses [11]. In the following studies, more than one scientific team demonstrated the possibility of PZQ-resistance [12–17]. All the evidence indicate that reliance on a single drug is not sustainable, searching for novel antischistosomal compounds is of priority for the treatment and control of schistosomiasis.

Unfortunately, despite the comprehensive use of PZQ, the mechanism of action and the targets of PZQ is still unknown, which exacerbates the difficulty of rationally designing potential drug candidates for schistosomiasis. Very few novel PZQ analogues and derivatives have been developed into clinical agents for the treatment of schistosomiasis japonica. In addition to PZQ, artemisinin and its derivatives show promising activity against juvenile schistosomes but have less efficacy against adult worms [18–19]. However, their use for schistosomiasis may be restricted in malaria-endemic areas to avoid putting its use as an antimalarial at risk. As reviewed by da Silva and coworkers [8], there are three strategies for development of new PZQ analogues: (a) synthesis of PZQ analogues; (b) the rational design of new pharmacophores; (c) the discovery of new compounds from screening programs on a large scale [8]. Owing to the unknowing mechanism of action and targets of PZQ, it is difficult to design new pharmacophores, the synthesis of new active analogues is possible by understanding the pharmacokinetics and chemical structure of PZQ. PZQ undergoes rapid metabolism and is converted into a major *trans*-cyclohexanol metabolite, which is much less effective than PZQ itself [20–21]. The ketone oxidation product of the *trans*-cyclohexanol metabolite and other analogues with increased metabolic stability were designed and had low to modest activity against juveniles of *S. japonicum* and *S. mansoni* [22–23]. These results imply that structural features of the cyclohexyl group (R group, Fig 1A) are likely related to antischistosomal activity. Several structural changes were made in the R position in order to increase the schistosomicidal activity. However, the vast majority of the derivatives demonstrated only low to moderate effect, their schistosomicidal activities were not comparable to PZQ [22–26]. In this study, we synthesized a new praziquantel derivative P96 with the cyclohexyl group substituted by cyclopentyl group (Fig 1B), with the aim of preventing the formation of 4-*trans*-cyclohexanol and increasing the metabolic stability. In this study, the derivative P96 was tested *in vitro* and *in vivo* against juvenile and adult stages of *S. japonicum*. Besides worm burden and egg burden, quantitative real-time PCR was employed to evaluate the antischistosomal efficacy at molecular level *in vivo*.

**Fig 1.**
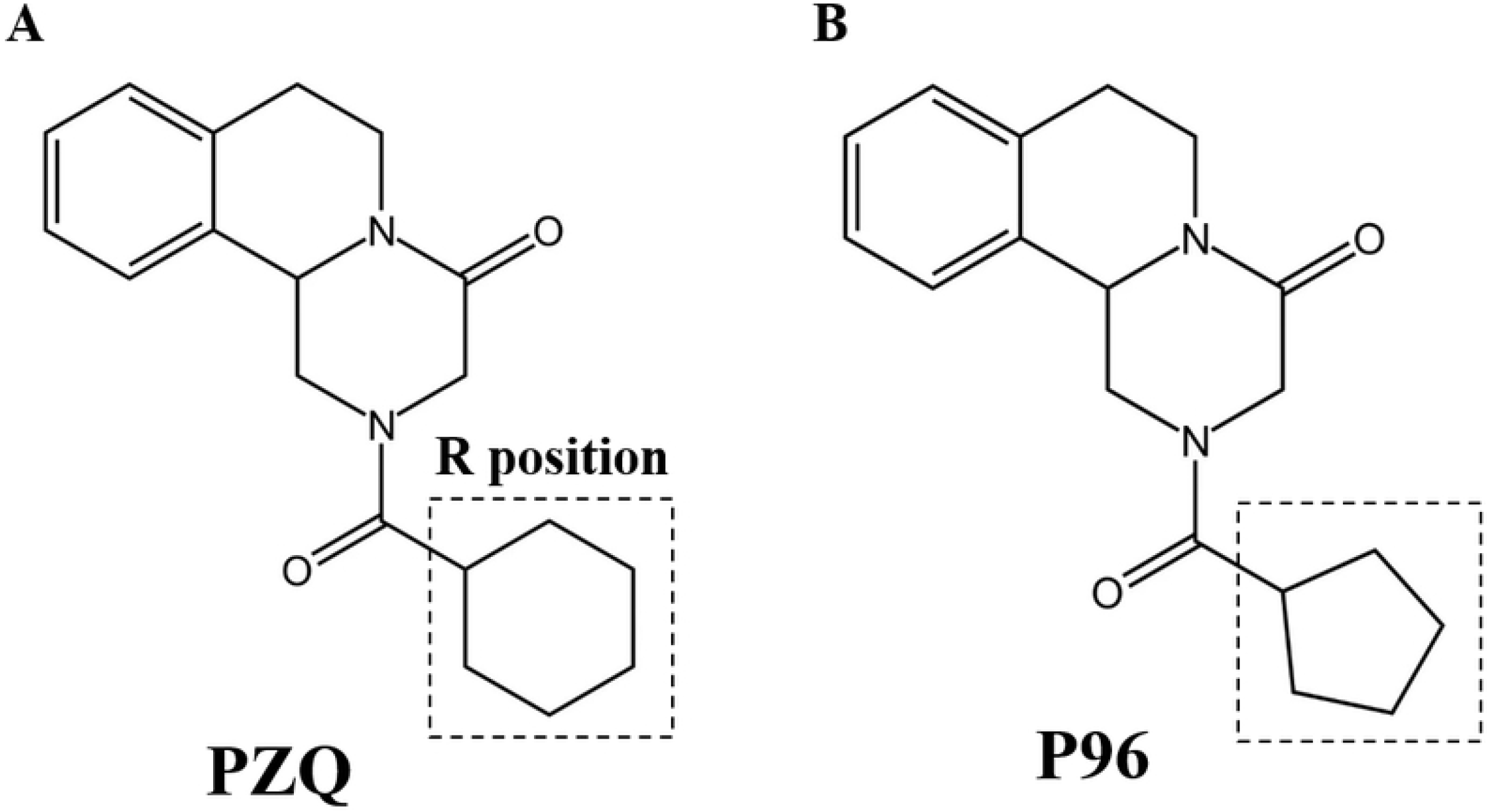
Chemical structures of praziquantel (PZQ) and P96.

## Materials and Methods

### Ethics statement

All the animal experiments were carried out in strict accordance with the recommendations in the Guide for the Care and Use of Laboratory Animals of the National Institutes of Health. The protocol (including mortality aspects) was approved by the Committee on the Ethics of Animal Experiments of the Soochow University (Permit Number: 2007-13).

### Parasites and animals

*S. japonicum* infected snail (*Oncomelania hupensis*) were provided by the Institute of Schistosomiasis Control in Jiangsu Province (Wuxi, China). *S. japonicum* cercariae (Chinese mainland strain) shedding from the snails were used to infect mice models. Female ICR mice (4-6 weeks-old and weighing 15-25g) and female New Zealand rabbits (weighting 2.0-2.5kg) were provided by the Experimental Animal Center of Soochow University (Suzhou, China). All mice and rabbits were raised under specific pathogen-free conditions with controlled temperature (20 ± 2 °C) and photoperiod (12 h light, 12 h dark). Each mouse was transcutaneously infected with 60±2 *S. japonicum* cercariae. Each rabbit was infected with 200±5 *S. japonicum* cercariae.

### Reagents

PZQ analogue P96 was synthesized by School of Pharmaceutical Sciences of Shandong University. PZQ powder was purchased from Sigma-Aldrich (St. Louis, MO, USA). Dulbecco’s modifed Eagle’s medium (DMEM) and penicillin/streptomycin were purchased from Life Technologies (Carlsbad, CA, USA). New-born calf serum was purchased from Biological Industries (Cromwell, CT, USA). *In vitro*, all chemicals were dissolved in dimethyl sulfoxide (DMSO, Fluka, Buchs, Switzerland). *In vivo*, all compounds were dissolved in corn oil.

### *In vitro* treatment

Worms recovered from *S. japonicum* infected mice at 16 days (juvenile worms) and 35 days (adult worms) post-infection were collected through perfusion of the hepatic portal system and mesenteric veins [27]. The worms were placed in 6-well plates (Corning Costar, Corning, New York, USA) containing Dulbecco’s modified minimum Eagle’s medium (bicarbonate buffered) supplemented with 10% newborn calf serum, 100 U /ml penicillin and 100 μg/ml streptomycin, and incubated at 37 °C in an atmosphere of 5% CO_2_ in air. Juvenile worms were divided into four groups, with five worms per well, each being tested in triplicate, as follows: group I, untreated control, incubated with complete DMEM containing 0.1% DMSO; group II, worms treated with 25 μM P96; group III, treated with 50μM P96; group IV, treated with 100μM PZQ. Adult worms separated by sex accepted the same treatment as juveniles. All the worms were exposed to the different compounds for about 16h, then washed three times with sterile saline, and subsequently cultured in drug-free medium. At 24, 48 and 72h post-incubation, the worms were observed under a dissecting microscope (SZX16, Olympus, Japan), and viability score was assigned as described previously [28], based on the changes of mobility and general appearance. Briefly, viability score of each worm ranged from 0 to 3: Worms with the highest score of 3, as observed in the control group during the observation period, moved more actively and softly, and the body was transparent; 2: Worms moved their entire bodies but stiffly and slowly, with the body translucent; 1: parasites moved partially and had an opaque appearance; 0 points: the worms remained contracted and did not resume movement, deemed as ‘dead’.

### Scanning electron microscopy (SEM)

Ultrastructural features of tegument of schistosomes treated with P96 and PZQ were examined using SEM and were compared with control group and PZQ treatment group. For SEM, the schistosomula and male adult worms were washed three times in phosphate-buffered saline (PBS; pH 7.4) and fixed overnight at 4 °C in 2.5% glutaraldehyde-PBS solution (pH 7.4). After fixation, the worms were washed again in PBS, post-fixed in 1% osmium tetroxide, dehydrated in graded ethanol, then dried for approximately 30 min. Finally, the samples were mounted on aluminum stubs, coated with gold, and examined under a Hitachi-S4700 scanning electron microscope (Chiyodaku, Japan).

### *In vivo* treatment in mice of schistosomiasis japonica

For understanding the effect of P96 on different developmental schistosomes *in vivo*, female ICR mice infected with 60 ± 2 *S. japonicum* cercaria were randomly divided into 15 groups, with 10 mice in each group. Group 1, untreated control group, received vehicle (corn oil) only. Group 2-8, treated with an oral dose of 200mg/kg P96 for 5 consecutive days at 1 day, 3 days, 7days, 14 days, 21 days, 28days and 35 days post-infections, respectively. Group 9-15, treated with a single dose of 200mg/kg PZQ at the same time schedule as treated with P96. In order to understand whether there was a dose-dependent effect of P96, *S. japonicum* infected mice were treated with P96 at a single oral dose of 100, 200, 400, 600 mg/kg for 5 consecutive days, with 8 mice in each group. At 21 days post-treatment, all mice were sacrificed to assess the worm burden and worm reduction rate.

### *In vivo* treatment in rabbits of schistosomiasis japonica

A total of 8 female New Zealand White rabbits, weighing approximately 2.0-2.5kg, were randomly divided into 4 groups of 2 rabbits each. Each rabbit was infected with 200 ± 5 *S. japonicum* cercaria. Group1, untreated control group, received vehicle (corn oil) only. Group2, treated orally with150mg/kg P96 at 28 days postinfection. Group3, treated orally with 300mg/kg P96 at 28 days postinfection. Group4, treated with a single oral dose of 150mg/kg PZQ. After 3 weeks posttreatment, all the rabbits were sacrificed to recover the adult worms separated by sex for measuring the real worm burden and worm reduction rate.

### Rabbit blood sample collection

Pre-infection blood samples of rabbits were collected as negative control before infection. For all the groups, *S. japonicum* infected blood samples were collected on the 3rd day and then weekly until 4 weeks postinfection. Blood from PZQ-treated and P96-treated rabbits were collected once a day in the first week, and then weekly until 6 weeks posttreatment. Serum of each blood sample was separated by centrifugation (2000g for 10 min) after storage at 37 °C for 1 h. The sera were stored at −20 °C until DNA extraction.

### DNA extraction

DNA from all the collected serum samples was extracted using the method described previously [29], with slight modifications. Briefly, 200 μL of infected rabbit serum were dissolved in 400 μL serum lysis buffer containing 150 mM NaCl, 10 mM EDTA, 10 mM Tris-HCl (pH 7.6), 2% SDS, 5 μg/mL salmon sperm DNA, and 250 μg/mL proteinase K (Takara, Dalian), incubated at 55 °C for 1h, then extracted twice with phenol-chloroform-isoamyl alcohol (25:24:1) and precipitated with dehydrated alcohol. The DNA pellet was air-dried and dissolved in 25 μL of TE buffer (10 mM Tris-HCl, 1 mM EDTA, pH 8.0).

### Design of primers

As shown in Table 1, primers were designed targeting SjR2 retrotranposons of *S. japonicum*. Probes were designed with 5’ terminal reporter dye FAM and 3’ terminal quencher dye TAMRA. The specificity of primers and probes were tested using a BLAST search against the Genbank database.

**Table 1.**
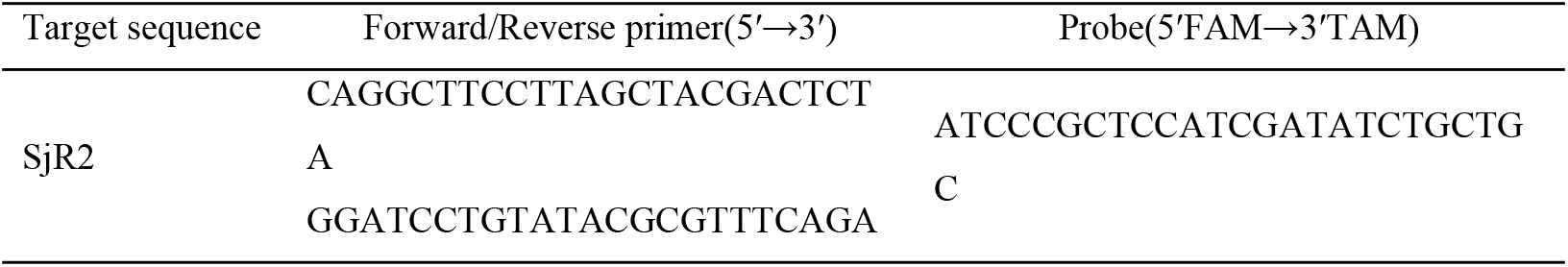
Primers and probes for SjR2 quantitative real-time PCR.

### Quantitative real-time PCR

The 25μL reaction mix contained 4μL DNA, 12.5μL 2×Platinum qPCR Supermix-UDG (Invitrogen by life technologies), 1μL 50mM MgCl_2_, 1μL ROX Reference Dye (1:10), 200nM of each primer, 100nM of probe, and distilled water to give the final volum of 25μL. The program consisted of incubation at 50°C for 2 minutes, followed by 95°C for 2 minutes, then 45 cycles at 95°C for 15 seconds and 60°C for 45 seconds. Ten-fold serial dilutions of standard plasmid with targeting sequence of SjR2 were used to generate the standard curve to calculate the copy numbers of SjR2 DNA (data not shown).

### Statistical Analysis

All data sets were analyzed using the SPSS26.0 software package. Data of viability score were expressed as the mean value ± standard error (SE). Data of worm number and egg burden were expressed as the mean value ± standard deviation (SD). Differences between groups were analyzed by one-way ANOVA followed by Dunnett’s test. Statistical significance of the difference of the sample rates was determined by the chi-square test. A *P*-value<0.05 was considered to be statistically significant.

## Results

### P96 exhibits potent schistosomicidal effect against both juvenile and adult worm *in vitro*

*In vitro*, after 24h exposure to different concentrations of P96 and PZQ, the mean viability score of males, females and juveniles was significantly decreased compared to the control group (males: *F*_(3,68)_=260.245, *P*<0.0001; females: *F*_(3,68)_=91.866, *P*<0.0001; juveniles: *F*_(3,68)_=86.821, *P*<0.0001;). The antischistosomal effect of P96 was concentration-dependent, with P96 at 50μM demonstrated the most obvious schistosomicidal effect against male, female and juvenile worms. The viability reduction rate of P96 at concentration of 50μM was 96.7%, 80% and 93.3%, respectively (Table 2), which was similar (females: *P*=1.000) as or even higher (males: *P*<0.05; juveniles: *P*<0.0001) than 100μM PZQ-treated group. Unlike PZQ, the lethal effect of P96 was not time-dependent. As shown in Fig 2, from 24 h to 72 h incubation period, the viability score of P96 treated group sustained at the same level.

**Fig 2.**
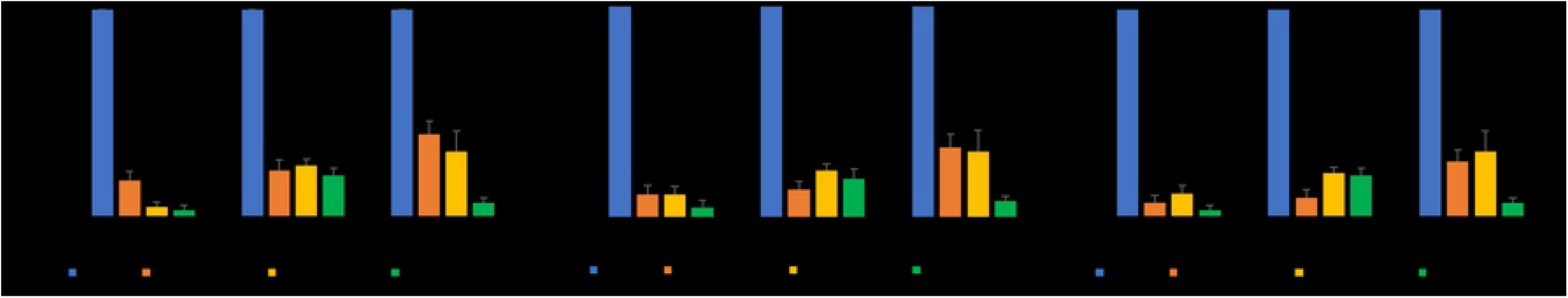
*In vitro* activity of P96 against males, females and juveniles of *S. japonicum*. Adult and juvenile worms were incubated with different concentrations of P96 and PZQ for 24h (A), 48h (B) and 72h (C). The viability was assigned using a viability score of 0-3. The control group was incubated with complete DMEM with 0.1% DMSO. ****represents significant differences compared to the control group, *P*<0.0001. ***represents significant differences compared to the PZQ treatment group, *P*<0.001. **represents significant differences compared to the PZQ treatment group, *P*<0.01. *represents significant differences compared to the PZQ treatment group, *P*<0.05.

**Table2.**
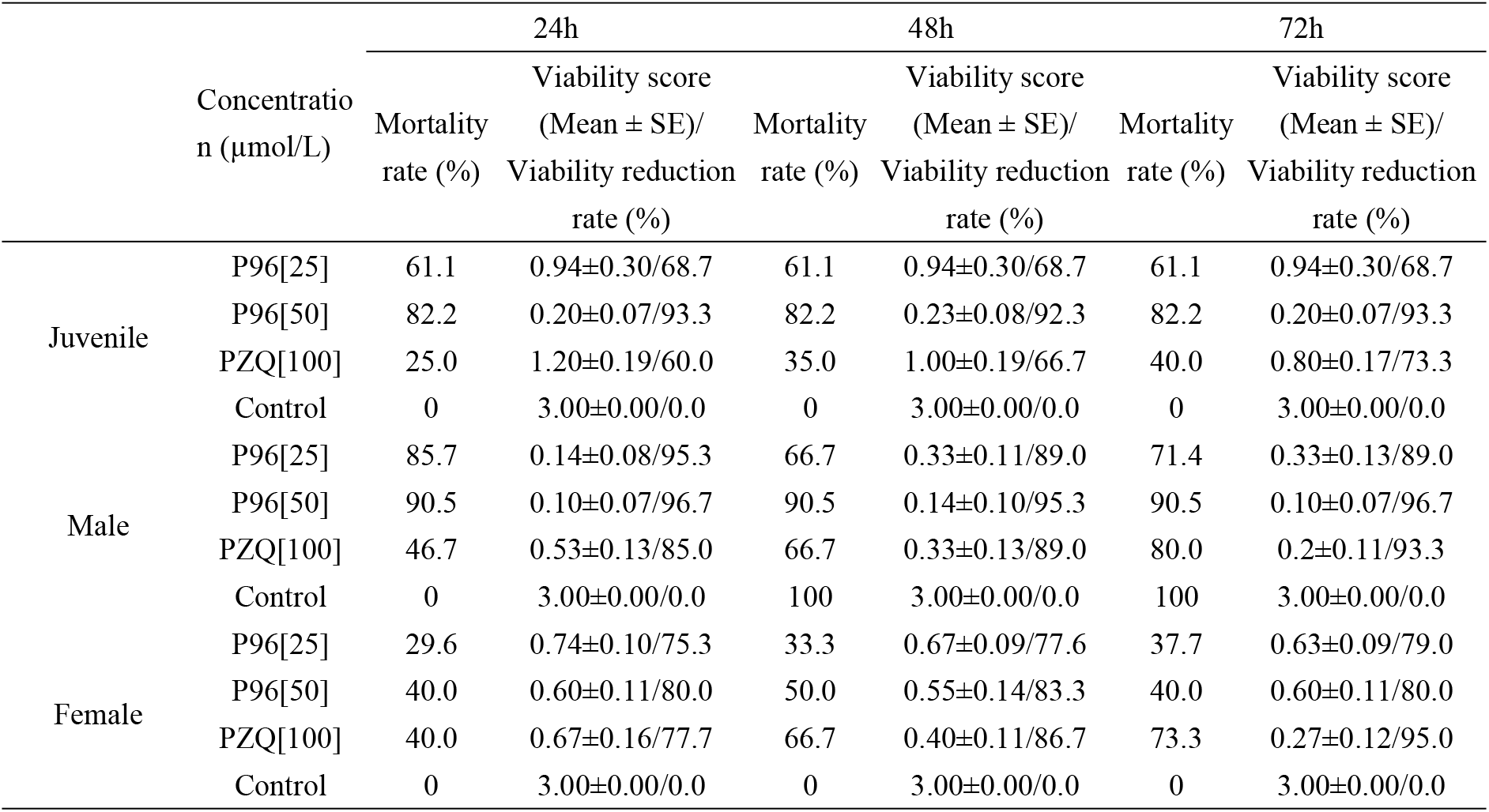
*In vitro* effect of P96 at different concentrations against juveniles, males and females of *Schistosoma japonicum*.

### Morphological properties by scanning electron microscopy (SEM)

SEM studies revealed that schistosomula from control group demonstrated normal tegumental ultrastructure features (Fig 3A). Numerous ridges were uniformly arranged along the mid-body of the schistosomula (Fig 3A). After treatment with 100 μM PZQ, the ridges became swollen, however, the integrity of the tegument was not compromised (Fig 3B). In contrast, the juveniles exposed to 50 μM P96 showed significant changes in the tegument. Extensive sloughing of the tegument and severe swelling were recorded (Fig 3C). Under SEM, male *S. japonicum* worms from control group showed normal tegument ultrastructures (Fig 4A). The tegument of the mid-body was intact, the crests with sensory papillae were uniformly arranged along the body (Fig 4A). The inner wall of gynecophoral canal and typical ridges were preserved (Fig 4B). Males exposed to 100 μM PZQ demonstrated disarrangement of crests with swelling sensory papillae in the tegument (Fig 4C). The ridges in the inner wall of gynecophoral canal were shallow or even disappeared. Pronounced oedema, collapsed papillae and shallow peeling were observed in this area (Fig 4D). Alterations in the tegument treated by 50μM P96 were different from that of PZQ. The tegumental structures were destroyed. The normal crests in the tegument of mid-body disappeared and fused into trabeculae. The sensory papillae were swollen, disformed, collapsed or even disappeared (Fig 4E). Severe swelling and extensive peeling of the tegument were detected in the gynecophoral canal inner wall (Fig 4F).

**Fig 3.**
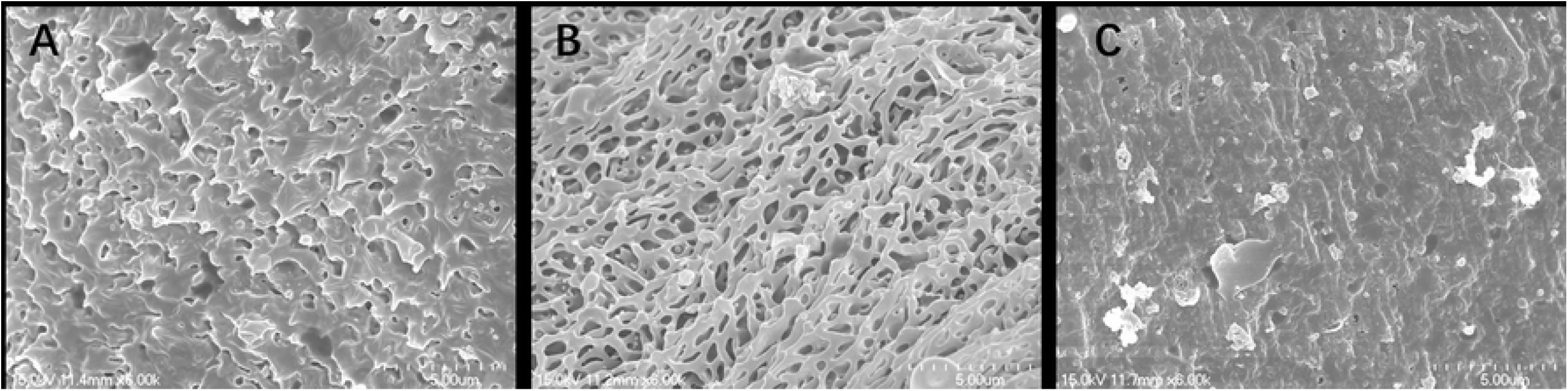
Scanning electron microscopy (SEM, × 3000) observation on the tegument of schistosomula. (A) mid-portion of the control schistosomula; (B) mid-portion of the worm exposed to 100μM PZQ; (C) mid-portion of the worm exposed to 50μM P96.

**Fig 4.**
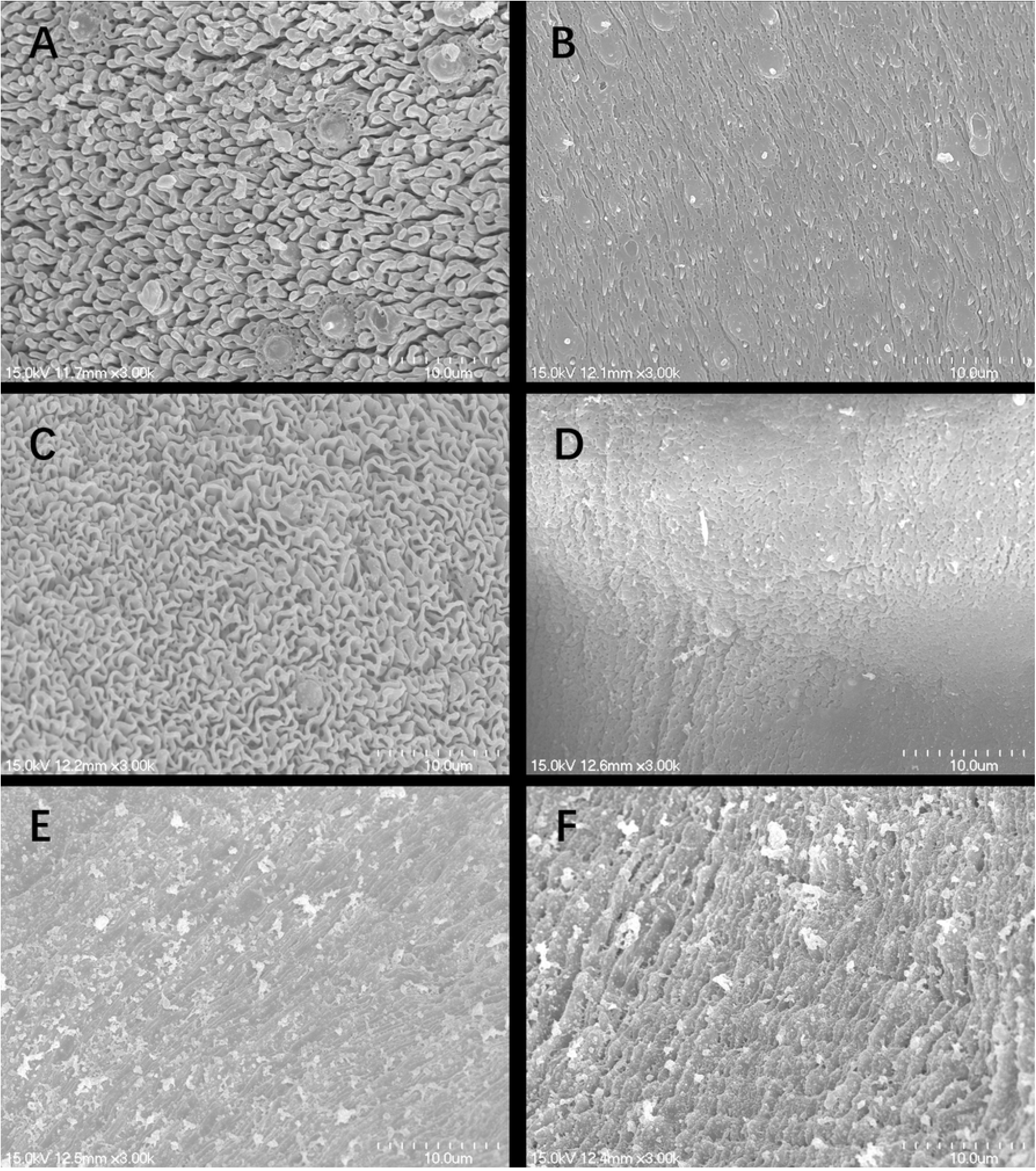
Scanning electron microscopy (SEM, × 3000) observation on the tegument of male adult *S. japonicum*. (A) mid-portion of the control worm; (B) inner wall of gynecophoral canal of control worm; (C) mid-portion of the worm exposed to 100μM PZQ; (D) inner wall of gynecophoral canal of the worm exposed to 100μM PZQ; (E) mid-portion of the worm exposed to 50μM P96; (F) inner wall of gynecophoral canal of the worm exposed to 50μM P96.

### Stage-sensitivity of P96 *in vivo*

As shown in Table3, the worm reduction rate caused by 200mg/kg P96 in mice harbored with1-day, 3-day, 7-day and 14-days juvenile of *S. japonicum* ranged from 43.5% to 58.2%, which was significantly higher than 200mg/kg PZQ treated group (9.0-27.5%), the *P* value was all lower than 0.05. On day 21, the effect of P96 was 45.9%, while 42.7% worm burden reduction for PZQ was observed. On day 28, adult worm stage, the worm reduction rate of P96 was 53.6%, which was similar to that of PZQ (67.1%, *P*=0.157>0.05). On day 35, adult worm pairing and spawning stage, PZQ exerted the most outstanding activity, the worm reduction rate was 96.1%, however, not significantly higher than the reduction of 86.9% for P96 (*P*=0.163>0.05).

**Table 3.** *In vivo* activity of P96 at a single oral dose of 200mg/kg for 5 consecutive days against different developmental stages of *S. japonicum* in mice.

|  | Worm number (mean±SD)/worm reduction(%) |  | Chi-square test | *P* value |
| --- | --- | --- | --- | --- |
|  | P96(200 mg/kg) | PZQ(200 mg/kg) |  |  |
| Control <sup>a</sup> | 51.0 ± 2.1/0.0 | 51.0 ± 2.1/0.0 | / | / |
| 1-day-pi | 23.5 ± 3.5/53.9 | 37.0 ± 1.9/27.5 | $\chi^2=7.428$ | 0.006 |
| 3-day-pi | 28.8 ± 4.6/43.5 | 46.3 ± 4.6/9.8 | $\chi^2=14.557$ | <0.0001 |
| 7-day-pi | 28.5 ± 3.5/44.2 | 46.4 ± 3.4/9.0 | $\chi^2=15.417$ | <0.0001 |
| 14-day-pi | 21.3 ± 3.2/58.2 | 41.3 ± 2.7/19.9 | $\chi^2=16.452$ | <0.0001 |
| 21-day-pi | 27.6 ± 5.0/45.9 | 29.2 ± 3.3/42.7 | $\chi^2=0.04$ | 0.842 |
| 28-day-pi | 23.6 ± 2.3/53.6 | 16.7 ± 2.9/67.1 | $\chi^2=1.998$ | 0.157 |
| 35-day-pi | 6.7±0.58/86.9 | 1.7±0.6/96.7 | $\chi^2=1.950$ | 0.163 |

<sup>a</sup>: Mice were given an equal volume of corn oil, pi: postinfection.

### The dose-response of P96 against *S. japonicum* juveniles in mice

Considering the promising antischistosomal efficacy of P96, we further assessed its effect against 14-day-old juveniles in *S. japonicum* infected mice. As shown in Table 4, with the ascending dose of P96 from 100 mg/kg to 600 mg/kg, the mean worm burden decreased, and the worm reduction rate increased from 48.1% to 68.4%, indicating that there was a dose-dependent effect of P96 against juveniles. Meanwhile, with the increasing dose of P96, the mortality of the mice declined remarkably. Fifty percent mice died in 100 mg/kg treated-group, 12.5% mice died in 200 mg/kg treated-group, no mice died in 400 mg/kg and 600 mg/kg treated-group.

**Table 4.**
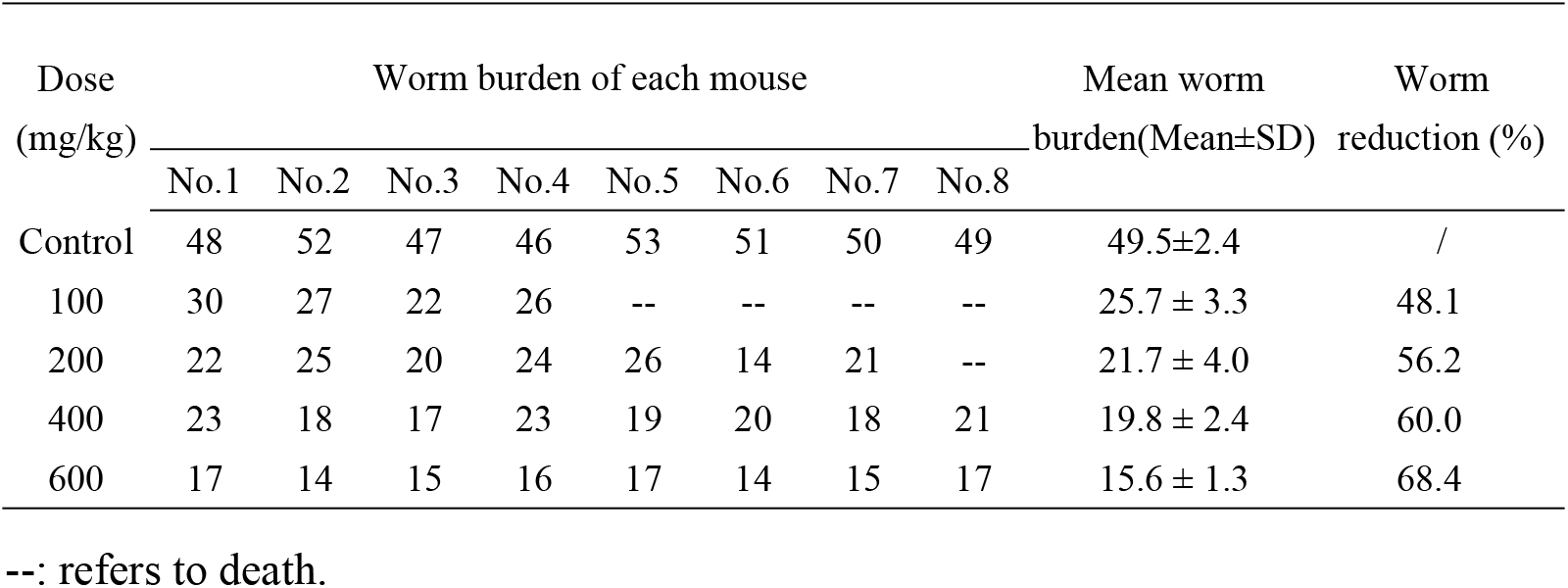
The dose-dependent effect of P96 against 14-day-old *Schistosoma japonicum* juveniles in mice with a daily oral dose 100-600 mg/kg for 5 days.

| Dose<br>(mg/kg) | Worm burden of each mouse |  |  |  |  |  |  |  | Mean worm<br>burden(Mean±SD) | Worm<br>reduction (%) |
| --- | --- | --- | --- | --- | --- | --- | --- | --- | --- | --- |
|  | No.1 | No.2 | No.3 | No.4 | No.5 | No.6 | No.7 | No.8 |  |  |
| Control | 48 | 52 | 47 | 46 | 53 | 51 | 50 | 49 | 49.5±2.4 | / |
| 100 | 30 | 27 | 22 | 26 | -- | -- | -- | -- | 25.7 ± 3.3 | 48.1 |
| 200 | 22 | 25 | 20 | 24 | 26 | 14 | 21 | -- | 21.7 ± 4.0 | 56.2 |
| 400 | 23 | 18 | 17 | 23 | 19 | 20 | 18 | 21 | 19.8 ± 2.4 | 60.0 |
| 600 | 17 | 14 | 15 | 16 | 17 | 14 | 15 | 17 | 15.6 ± 1.3 | 68.4 |

--: refers to death.

### The dose-response of P96 against *S. japonicum* adults in rabbits

Table 5 summarized the activity of P96 at different doses against 28-day-old adult worm in *S. japonicum* infected rabbits. Rabbits treated with 150 mg/kg P96, resulted in a statistically significant reduction in the mean total worm burden and egg burden compared with the control group (worm burden: *F*_(3,6)_=139.655, *P*<0.0001, egg burden: *F*_(3,50)_=47.399, *P*<0.0001). The worm reduction rate and the egg reduction rate were 65.2% and 80.1%. With the dose of P96 increasing to 300 mg/kg, the worm reduction rate was enhanced to 91.7%, which was very close to that of PZQ (98.5%) at the dose of 150 mg/kg (*P*=0.661>0.05).

**Table 5.** Effect of P96 at different oral doses against 28-day-old *Schistosoma japonicum* adults in rabbit models.

| Compounds | Worm burden (Mean±SD) |  |  | Worm reduction rate(%) | Egg burden/g liver tissue (Mean±SD) | Egg reduction rate(%) |
| --- | --- | --- | --- | --- | --- | --- |
|  | total | male | female |  |  |  |
| P96 (150mg/kg) | 58.5±11.2<br>* | 41.8±11.0 | 16.8±8.3 | 65.2 | 3257.8±1618.9* | 80.1 |
| P96 (300mg/kg) | 14.0±9.9 | 10.5±7.8 | 3.5±2.1 | 91.7 | 812.9±463.1 | 95.0 |
| PZQ (150mg/kg) | 2.5±0.7 | 1.5±0.7 | 1.0±0.0 | 98.5 | 400.0±228.9 | 97.6 |
| Control (corn oil) | 168.0±4.2 | 90.0±1.4 | 78.0±2.8 | / | 16378.3±8836.6 | / |

\* $P<0.0001$ , significant difference compared with control group.

As shown in Fig 5, among the three groups of rabbits treated by different doses of PZQ and P96 at 28 days postinfection (adult worm stage), the detection results of quantitative real-time PCR showed that the copies of SjR2 DNA were the highest at 3 days posttreatment of 150mg/kg PZQ (Fig 5), which was consistent with the fact that PZQ was the most effective against schistosome adult worm. In rabbits with chemotherapy of P96 at an oral dose of 150mg/kg, the content of SjR2 DNA also reached the peak on the 3^rd^ day posttreatment, but lower than PZQ treatment group (Fig 5). With the oral dose of P96 increasing to 300mg/kg, the peak of SjR2 DNA after 3 days posttreatment was higher than that of 150mg/kg P96 treatment group, indicating that there was a dose-dependent effect of P96 against schistosome adult worm. Moreover, the content of SjR2 copies in sera of rabbit with administration of 300mg/kg P96 was close to that of 150mg/kg PZQ treatment group (Fig 5).

**Fig 5.**
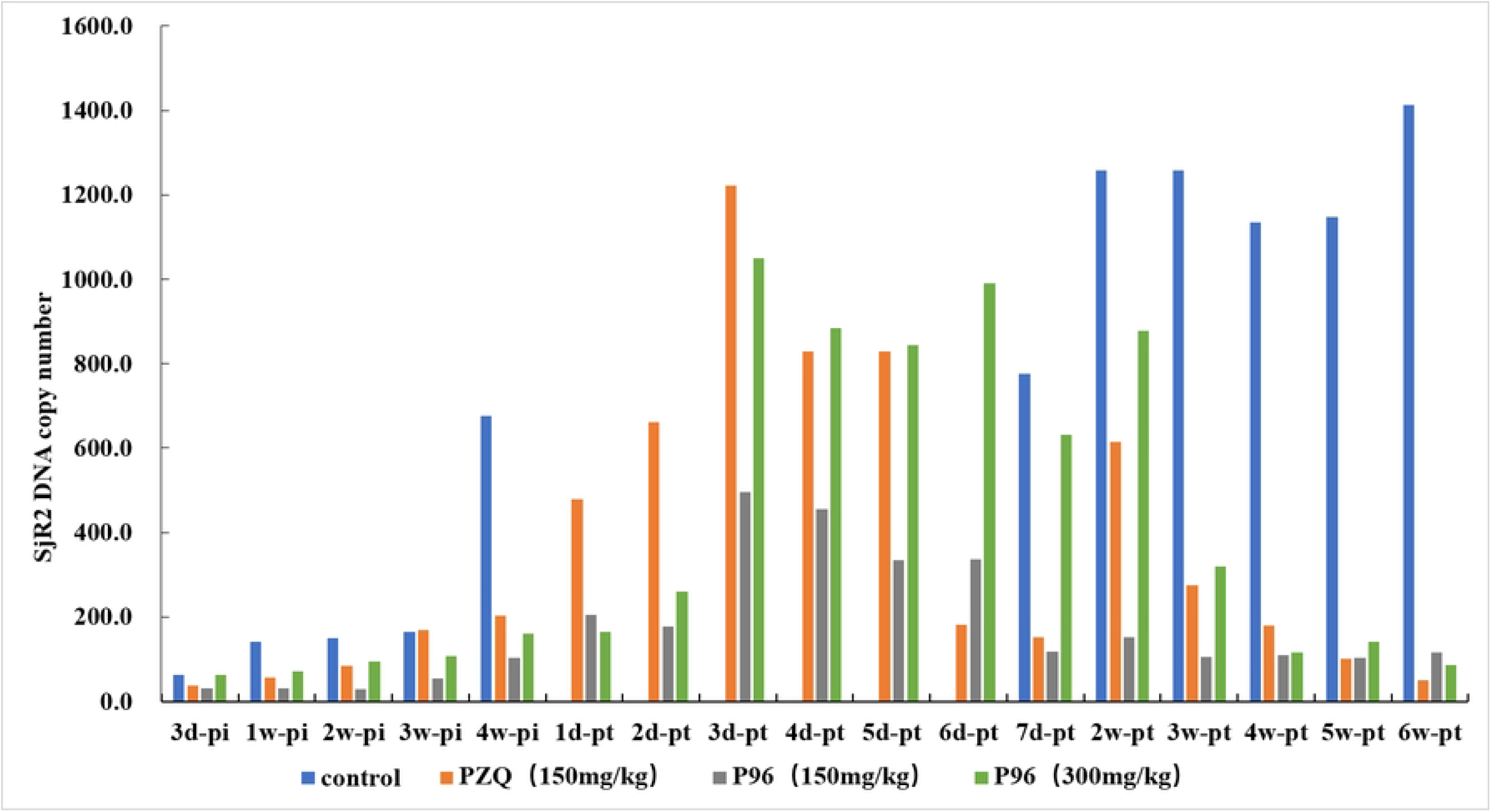
Dynamic changes of SjR2 DNA in sera of rabbits of schistosomiasis after chemotherapy of P96.

## Discussion

The large-scale and mono-therapeutic use of PZQ has raised many concerns for the treatment and control of schistosomiasis. The concerns mainly focused on the lack of activity against immature schistosome and the growing inclination to resistance of PZQ, which could be explanations to the poor cure rates and treatment failures in residents of high-risk regions [30]. Thus, it is of increasing importance to develop new drugs in the face of potential resistance to PZQ for the treatment of schistosomiasis, and the absence of an effective vaccine [31]. Even though PZQ has been the first treatment for several decades, the exact mechanism of action of the drug is still poorly understood. Hence, it is difficult to derive new compounds focusing on the same molecular target. However, synthesis of new PZQ derivatives might be a good strategy to develop novel antischistosomal agents [32]. Substantial research has been conducted to the design and development of PZQ derivatives. The chemical modifications are mainly concentrated on the cyclohexane ring and aromatic ring of praziquantel [33–34]. Unfortunately, vast majority of the analogues are not promising compounds, with low or moderate antischistosomal activity, being not comparable to PZQ [22, 26, 35–36]. Although no compelling agents have gone to clinical testing and trials, the results are of major importance for the analysis of its chemical structure for its activity of PZQ. As mentioned by Patra et al, whose studies demenstrated that the C10 aromatic ring of PZQ is not suitable for structural modification [35]. To impede the metabolism and increase metabolic stability, three different cycloalkyls substituents (cyclobutyl, cyclopentyl and cyclohexyl) with a carbonyl group in its structure were used for the synthesis of the derivatives. The results exhibited as the size of ring with carbonyl group enlarged, antischistosomal activity of R-isomers increased from 41.0% to 60.0%, indicating the size of the ring is important for the activity, with the cyclohexyl has the most effective activity [22]. The available results indicated that the derivatives structurally linked to PZQ through the metabolically liable cyclohexyl ring position might not afford active derivatives [22].

With this information, a novel derivative P96 with a small change, substitution of cyclohexyl with cyclopentyl, was synthesized in our study. Interestingly, it was this small change that brought us a big difference. Unlike previously reported derivatives, most of them demonstrating less activity than PZQ, the novel derivative P96 exerted potent antischistosmal efficacy against both juveniles and adults of *S. japonicum in vitro*. The schistosomicidal effect of P96 *in vitro* was dose-dependent, with a concentration at 50μM demonstrating the best activity. Male adult worms seemed to be more sensitive than females of the same age (Table 2). As we know that PZQ is highly effective against adults but has poor activity against juveniles. However, in our current study, P96 exhibited the most prominent activity against 16-day-old juveniles (82.2% of mortality, Table 2) compared to PZQ (25.0% of mortality, *P*<0.0001, Fig 2), and still retained high efficacy against adult worms, with a significant reduction of 96.7% in male worm viability compared to 60% of PZQ after 24h exposure (*P*<0.05, Fig 2).

Despite P96 had a structure very similar to that of PZQ, it displayed obviously different activity characteristics compared with PZQ, especially against juvenile stage worms. It is well known that integrality of the tegument plays a key role for worm survival, the parasite is vulnerable to the host immune system because of the surface antigens exposure [8, 32]. Our SEM observation revealed that P96 caused severe damage to the schistosomula tegument, including sloughing of the tegument with a disordered surface, as reported in the *in vitro* study [23]. Whereas PZQ only caused very light damage to the juvenile tegument, the morphological change might partially explain the poor activity of PZQ against immature schistosomes. The ultrastructural alterations of male adult worm treated by P96 were similar to that of PZQ, indicating that the potent antischistosomal activities of P96 might be correlated with its effects on worm tegument.

Although our *in vitro* results showed that P96 had promising antischistosomal activity, it was worth noting that potential action *in vitro* did not translate to impressive killing *in vivo*. As reported by Patra et al [37], upon alteration of the organometallic moiety to Cr(CO)3, the derivatives exhibited marked activity against *S. mansoni in vitro*, however, they exerted low activity *in vivo* [37]. In this study, two kinds of animal models were employed to evaluate the efficacy of P96 *in vivo*. In *S. japonicum* infected mouse model, P96 demonstrated outstanding antichistosomal activity against all worm developmental stage. Specifically, it presented remarkable schistosomicidal activity against young stages superior to PZQ. Moreover, it retained the high efficacy against the 35-day-old adult *S. japonicum* with a reduction rate of 84.4%, which was similar to PZQ (Table 3). Our *in vivo* results also confirmed that the efficacy of PZQ was stage-dependent, as the worm getting mature, demonstrating the most activity (Table 3). In addition, the dose-response study in mice revealed that there was a dose-dependent effect of P96 against juveniles, with the highest oral dose at 600mg/kg achieved the maximum worm reduction rate (68.4%, Table 4). Thereafter, the activity of P96 was further tested in a large size rabbit model of schistosomiasis. The results also demonstrated a dose-dependent effect of P96 against adult worm. With the ascending dose of P96, the more promising reduction of worm burden was observed. At concentration of 300mg/kg, the worm reduction rate reached 91.7%, which was very close to 98.5% of PZQ (Table 5). However, further experiments are needed to identify the optimal dose for P96 *in vivo*.

Our quantitative real-time PCR detection results confirmed the prominent antischistosomal activity of P96. In our previous study, we have demonstrated that the specific SjR2 DNA of *S. japonicum* in rabbit sera mainly came from the residual body of dead worms and the disintegration of inactive eggs after chemotherapy of PZQ [38]. In this study, the SjR2 DNA detection results revealed that the content of SjR2 DNA achieved at the highest level on the 3^rd^ day posttreatment of 150mg/kg PZQ, indicating the higher copies of SjR2 DNA might represent better worm killing effect. In sera of rabbits with oral administration of 150mg/kg P96, the content of SjR2 DNA also reached its peak at 3 days posttreatment, but lower than that of PZQ treatment. With the dose of P96 increasing to 300mg/kg, the copy numbers of SjR2 DNA were close to 150mg/kg PZQ after 3 days treatment, which further verified that P96 has comparable activity to PZQ when tested against adult *S. japonicum*.

## Conclusion

Our *in vitro* and *in vivo* results demonstrated that P96, the novel PZQ derivative with a small structure change, cyclohexyl substituted by cyclopentyl, had broad-spectrum antischistosomal activity, especially against immature stages. Its remarkable schistosomicidal efficacy against both young stage and adult worm of *S. japonicum* enabled it could serve as the promising drug candidate for treatment and control of schistosomiasis. Further studies were needed to elucidate the *in vivo* metabolism of P96, as well as its mechanism of action against schistosomes.

## Acknowledgments

The authors would like to thank the Marine College of Shandong University for synthesis of the compound P96 for this study.

## Authors’ contributions

JX, DS and CX conceived and designed the study. LD, and HS acquired the experimental data. PH, RZ and XW contributed to data analyses. JX wrote the manuscript. All authors read and approved the final manuscript.

